# Convergent Evolutionary Traces of Genomic Innovations and Depletions in Socially Parasitic Ants

**DOI:** 10.1101/2024.12.31.630940

**Authors:** Mark C Harrison, Alice Séguret, Christopher Finke, Evelien Jongepier, Anna Grandchamp, Marah Stoldt, Jürgen Heinze, Susanne Foitzik, Erich Bornberg-Bauer, Barbara Feldmeyer

## Abstract

Parasitism, a common strategy across life, often involves genomic reduction. Socially parasitic “slavemaking” ants provide an excellent model to study the evolution of this lifestyle. Here, we compared the genomes of four closely related, independently evolved parasite-host pairs and their outgroups to identify convergent patterns. While genome size and gene numbers remain relatively stable, parasites show increased positive and relaxed selection, more putative *de novo* genes which have recently emerged from previously non-coding regions, and a significant loss of chemical receptors, particularly those associated with the evolution of sociality. Hosts, however, retain many duplicated genes. Gene co-expression networks associated with nest-defence in hosts are rather conserved, while the network structure of raiding parasites is much more variable and reflects the independent evolutionary origin of social parasitism. Our findings suggest that while hosts primarily rely on existing molecular mechanisms for defence, parasite genome evolution is characterised by extensive, convergent gene loss, innovation, and network rewiring.

## INTRODUCTION

Parasitism is a widely distributed lifestyle and the co-evolution between hosts and parasites is considered a major driver of evolution across many genera [Ebert and Fields, 2020]. Parasitism is often associated with genomic reductions [Sakharkar et al., 2004, Wernegreen, 2005, Schrader et al., 2021, Xu et al., 2021] which renders parasites irreversibly dependent on the host’s resources. A widely heralded host-parasite association is the exploitation of ant colonies through other, often closely related ants [Beibl et al., 2005, Prebus, 2017]. The social parasitism of ants includes inquilism, in which the parasite queen invades a host colony and exploits the workers there. Inquiline species have lost the worker caste and produce only sexuals as offspring. In contrast, “slavemaking” ants (also called “dulotic”, Greek for enslavement) have retained the worker caste; however, these do not exhibit typical worker behaviour such as brood care or foraging but have instead evolved a novel raiding behaviour. They specialize in raiding host colonies to abduct their worker brood, whose social behaviour they later exploit as adults [D’Ettorre and Heinze, 2001, Heinze et al., 2015].

Using comparative evolutionary genomics, both classes of parasitic ants were found to show signatures of genomic erosions and loss of chemoreceptors [Schrader et al., 2021, Jongepier et al., 2022]. Most prominently, odorant receptors (ORs), which are essential for sensing food and pheromones (cuticular hydrocarbons, CHCs) as well as identifying nest-mates and mating partners, were lost in both groups of socially parasitic ants [Schrader et al., 2021, Jongepier et al., 2022]. Numbers of gustatory receptors (GRs), which are essential for foraging, decreased even more in dulotic species [Jongepier et al., 2022]. Together with signals of relaxed selective constraints in GRs and ORs, which are both related to social organisation, these losses corroborate the reduced social behavioural repertoires in these parasitic ants. While acquired genes and their roles have been extensively studied, investigating the benefits of gene losses remains a challenge [Monroe et al., 2021]. Losses may, e.g., in response to environmental stimuli, render certain genes non-essential, release constraints, modify traits, or reduce metabolic costs and eventually lead to alternative evolutionary paths and increased fitness in specific environments. Thanks to the availability of many high quality ant genomes spanning short and long evolutionary time scales and computational analyses it is possible to reconstruct ancestral states and the courses of molecular changes leading to extant host and parasite species. Moreover, ant genomes are known for their rapid evolution, e.g., in the evolution of gene families, and their variability in gene composition between species [Simola et al., 2013]. The ant genus *Temnothorax* and its relatives offer a unique setting because nine host-parasite pairs evolved over a relatively short evolutionary period [Beibl et al., 2005, Prebus, 2017]. Comparative genomic analyses between these pairs can therefore aid in discerning stochastic from adaptive genomic changes, including losses, shedding light on their adaptive significance in driving evolutionary shifts and disentangling lineage-specific from recurrent events in the course of host and parasite evolution [Prebus, 2017].

Following on from a recent study [Jongepier et al., 2022] in which we found ORs to be convergently lost in three *Temnothorax* parasites, we here employ a broad comparative genomic approach. We identify signals of frequent gains or losses in protein families and selection signals among 18 ant species comprising one additional host-parasite pair, i.e., across four host-parasite pairs in total. Generalising the OR losses, we find that parasites lost many more genes than hosts. At the same time, parasites accumulate more lineage-specific putative *de novo* genes than their hosts, while the latter accumulate more duplicated genes. We find that parasites have substantially more protein families under positive and also under relaxed selection than hosts and that signals of selection relate primarily to difference in lifestyle. Revisiting the case of convergently lost ORs, we confirm that these losses affected mainly those which emerged in the early evolutionary history of ants and were likely crucial for their emerging social behaviour, which is partially lost in the social parasites. Finally, we ask if gene co-expression networks (GNWs) provide any indication of regulatory changes that have accompanied the evolution of parasites from a common ancestor with their hosts. When comparing one host and its parasite in a tandem running context (one ant leads at least one other ant to a food or nest source) we find striking similarities in the structure of the tandem-run GNWs, suggesting common evolutionary ancestry. However, comparing the GNWs within two hosts and within two parasitic species during a raid and outside raiding season we find networks in the raiding context underwent drastic changes.

## RESULTS

### Assembly and annotation of ant genomes

In a former study [Jongepier et al., 2022] we sequenced the genomes of three parasite ant species (*Harpagoxenus sublaevis*, *T. ravouxi* - formerly *Myrmoxenus ravouxi* -, *T. americanus* - formerly *Protomognathus americanus*), three host species (*Leptothorax acervorum*, *T. unifasciatus*, *T. longispinosus*), and two non-host sister species (*T. rugatulus*, *T. nylanderi*). In the present study, we sequenced one additional parasite genome, *T. pilagens*, and the genome of its main host, *T. ambiguus*. Collection, sequencing, assembly, and annotation followed our earlier study [Jongepier et al., 2022] (see Tables S1, S2, S3 for details on sample collection, sequencing, and genome assembly). These four host-parasite pairs and the two non-host species (*T. rugatulus*and *T. nylanderi*) form the *core set* of ten ant species that we analysed in this study. We used the genomes of five further ant species as *outgroups* to this core set, forming an *extended set* comprising 15 ant genomes. These included *Pristomyrmex punctatus*, *Myrmecina graminicola*, and *Acanthomyrmex ferox*, from the GAGA project [Boomsma et al., 2017], as well as the publicly available *Monomorium pharaonis* and *Cardiocondyla obscurior* genomes.

### Genes are more often convergently lost among parasites than hosts

We performed a gene family evolution analysis with CAFE5 [De Bie et al., 2006] to understand which gene families have expanded or contracted and if any of these changes may relate to the changes in lifestyles from non-host to either host or parasite. Overall, the numbers of significant gene family expansions was higher in host species (557 unique gene families) than in parasite species (455), while contractions were more abundant in parasite lineages (362) than in hosts (301) (Fig. 1A and Fig. 2A; Supplementary dataset 1).However, these differences were neither significant when controlling for phylogenetic relationships (expansions: F=0.738, p=0.278; contractions: F=0.494, p=0.337; phylogenetic ANOVA), nor when comparing against 1000 sampled pairs of four genomes which were randomly drawn from the extended set. However, differences between host and parasite species are more pronounced when only considering gene families convergently contracted and expanded in at least three out of four species (Fig. 2B). Among the extended set, when considering all 1365 combinations of four species, a median of five gene families were convergently, and significantly contracted among at least three out of four species. Among the four parasite species, 17 gene families were convergently contracted in at least three species, which corresponds to the 94*^th^* percentile (Fig. 2B). Six of these convergently contracted gene families were chemoreceptor (CR) families (5 GR and 1 OR), possibly linked to the loss of typical worker behaviours such as brood care and foraging in the parasites. Four further convergently contracted gene families have likely functions in pathogen resistance (see Supplementary dataset 1 & 2a), indicating that the loss of typical worker behaviours is linked to reduced pathogen exposure or that these parasites rely on their hosts’ immunity services, provided through social immunity behaviours and substances. In contrast, only three gene families were significantly contracted among at least three of the four host species (32*^nd^* percentile). Convergently expanded gene families were more numerous in hosts (13; 67*^th^* percentile), spanning various functions (Supplementary dataset 2b), than in parasite species (7 convergent extractions; 39*^th^* percentile; median across all species: 9; Fig. 2B).

**Figure 1:**
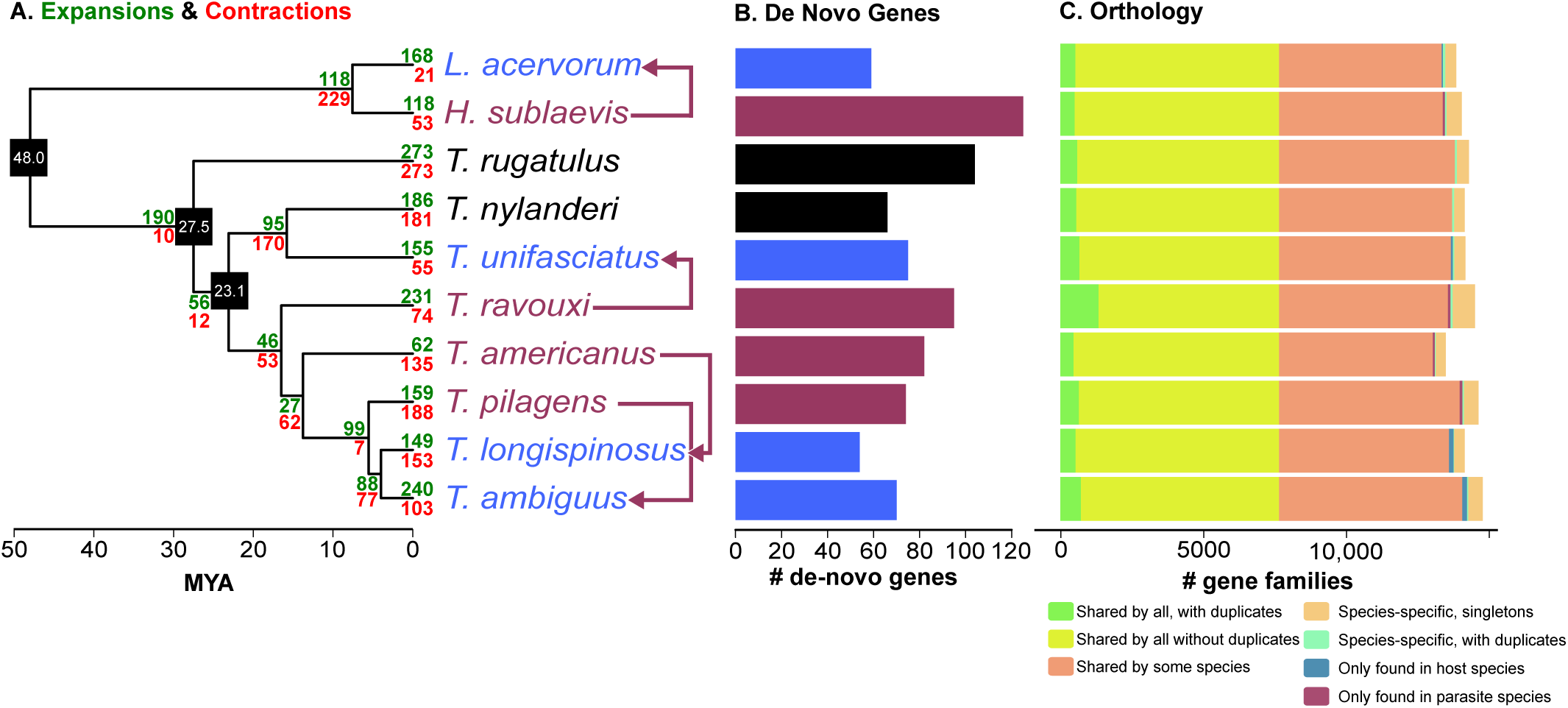
Proteome evolution of the core species set. A: Inferred numbers of significantly expanded and contracted gene families are shown in green and red, respectively. The three black nodes denote the anchors used for dating the tree with age given in the boxes. Species names are indicated on the branch tips according to lifestyles with parasites in magenta and hosts in blue, with arrows indicating parasite-host pairs, black indicating free-living non-host species; B: numbers of lineage specific putative (*de novo*) genes in each species, coloured by lifestyle (see also next). C: Numbers of inferred orthologous gene families. “only found in host (parasite) species” includes both, lineage specific putative de novo genes and those with orthologs only in other hosts (parasites). Those which have further evidence with an orthologous region found in synteny in a non-coding region in some outgroup genomes (see also Fig. S1) are the ones shown in (B) but at a different scale.

**Figure 2:**
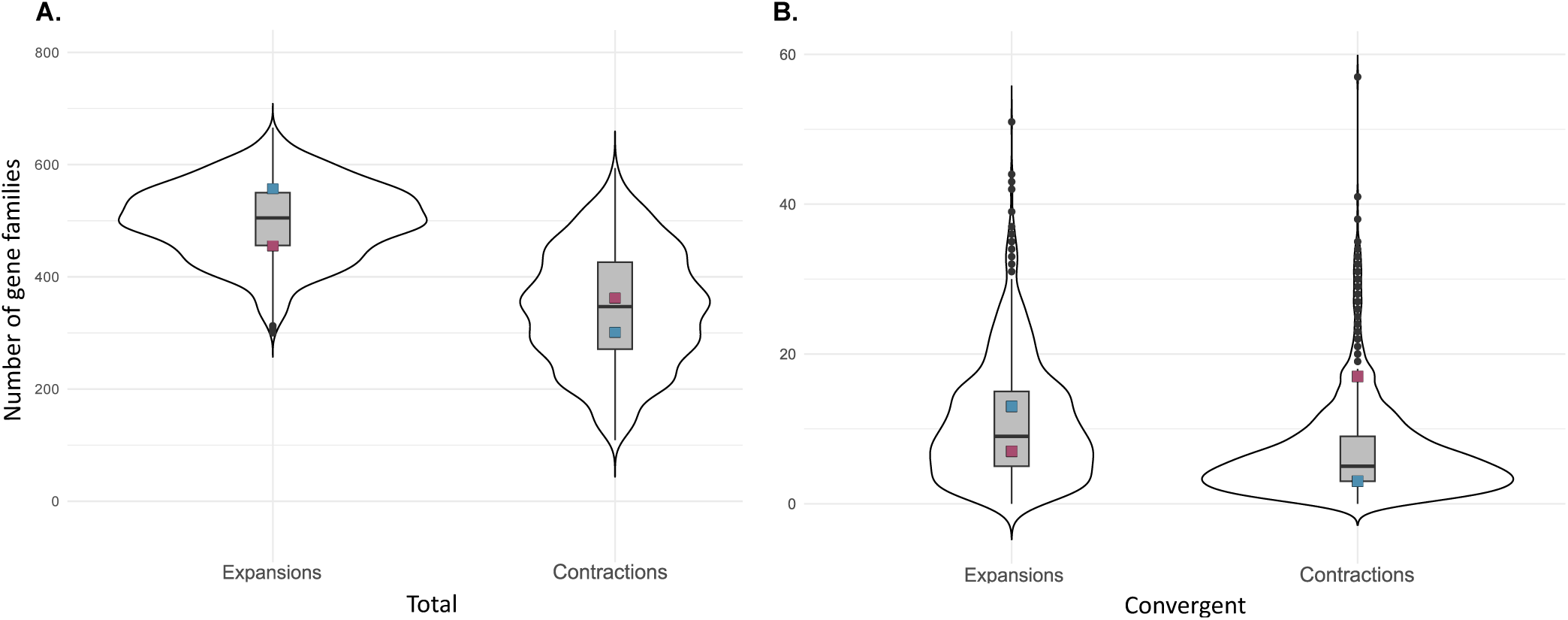
Convergent expansions and contractions. Shown are the numbers of gene families that are significantly expanded or contracted in all combinations of four species (n=1365 combinations of four out of the 15 analysed species in the extended set), with the number in the four host and parasite species shown by the blue and red boxes, respectively. A. shows the complete numbers of expansions and contractions within at least one of the four species. B. shows numbers of gene families convergently expanded and contracted in at least three out of four species.

### Convergently lost ORs emerged at the root of ants, likely related to the evolution of sociality

We manually annotated 5004 chemosensory receptor (CR) genes across all ten species of the core set, including 4030 OR and 974 GR genes. Confirming previous findings of convergent losses of ORs and GRs in three origins of social parasitism [Jongepier et al., 2022], we find both CRs to be significantly reduced in all four analysed parasite species compared to host and free-living species (ORs: F(2,7)=95.2, p=0.001; GRs: F(2,7)=18.5, p=0.006; phylogenetic ANOVA; Figs. 3A and S2A).

**Figure 3:**
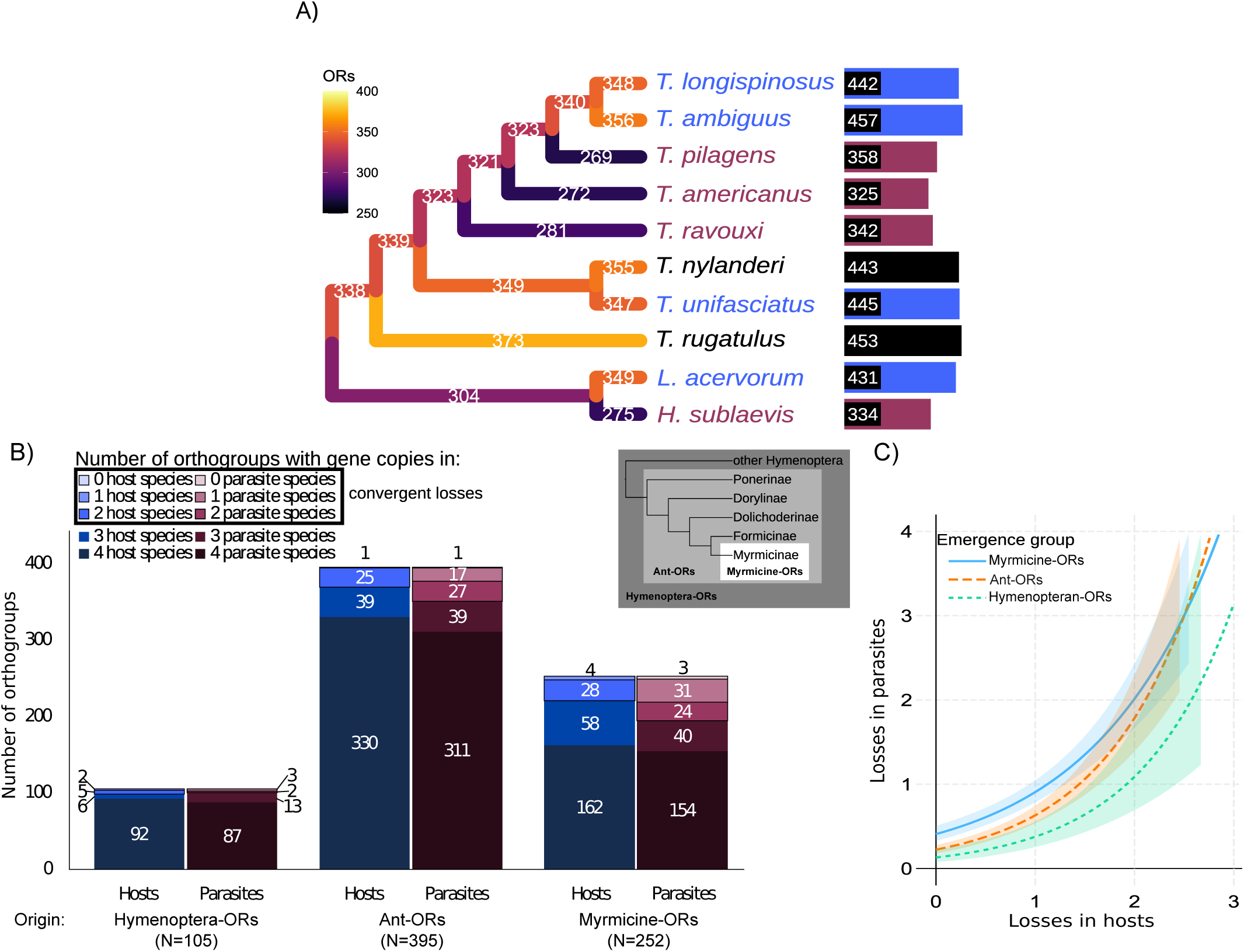
Evolution of odorant receptors. A) Ancestral numbers of ORs were reconstructed with CAFE5, with branches coloured by estimated counts. Counts were based on orthogroups containing at least 75% ORs. Numbers in black boxes show total numbers of ORs, including species-specific copies. B) Orthogroups of ORs are classified based on their predicted origin: “Myrmicine”-orthogroups only contain myrmicine-specific ORs. “Ants”-orthogroups only contain ant-specific ORs, including outside of Myrmicinae, such as: *Odontomachus brunneus*, *Harpegnathos saltator*, *Dinoponera quadriceps* (all Ponerine); *Eciton burchellii* (Dorylinae); *Linepithema humile* (Dolichoderine); *Camponotus floridanus*, *Formica selysi*, *Lasius fuliginosus*) (all Formicinae). “Hymenopteran” orthogroups contain ORs in species outside of ants: (*Vespula vulagaris*, *Nasonia vitripennis*, and *Athalia rosae*). Each group of orthogroups is again sub-classified based on how many have been lost in 0, 1, 2, 3 or all host species (blue) or parasite species (red). For example, there are 395 ant-specific orthogroups, of which 311 contain ORs in all 4 parasite lineages, whereas 39, 27, 17 and 1 ant-specific orthogroups have been lost in 1, 2, 3 and 4 parasite species, respectively. C) The result of a GLM that relates losses in host species to parasite species depending on the origin of the ORs. In all cases, convergent losses (in 2 or more species) are more prevalent in parasite species than in host species. However, the difference is significantly higher in ant-specific ORs (orange curve). GLM function used: glm(ParasiteLosses ∼ Origin * HostLosses, family = poisson).

We next investigated whether evolutionarily novel CRs are more frequently lost within parasite species than conserved CRs. For this, we established a *hymenopteran set* by supplementing the *extended set* with 11 further hymenopteran genomes, including another 8 ant genomes from other ant subfamilies and three non-ant hymenopterans (see Fig. 3).

We investigated orthology across all ORs and GRs with CAFE5 [De Bie et al., 2006]. We found that most of the 752 orthogroups of ORs emerged within ants outside of Myrmicinae (Ant-ORs: 395), while 252 having emerged within Myrmicinae (Myrmicine-ORs). 105 orthogroups also exist outside of ants, indicating an even older, pre-ant hymenopteran origin (Hymenotpera-ORs). Overall, most gene losses occurred among Myrmicine-ORs. Of these 252 relatively young Myrmicine-ORs 98 (38.9%) had been lost in at least one parasite species and 90 (35.7%) had been lost in at least one host species. This compares to relatively fewer losses in the older categories of Ant-ORs (12.4%) and Hymenopteran-ORs (21.3 %). When comparing between lifestyles, we measured greater losses of ORs within parasites than in hosts, regardless of their age (Fig. 3B). However, convergent loss (i.e. within at least 2 species) was significantly more common among parasites than hosts for Ant-ORs than for Myrmicine-ORs. This was evidenced by a significantly steeper slope relating losses in host species to losses in parasite species for Ant-ORs (p=0.0315; GLM, poisson distribution; Fig. 3C). Overall, this suggests that OR losses are more likely to manifest in parasites, with convergent losses particularly involving ORs which have emerged in the early evolutionary history of ants, during a time when ants evolved their highly complex social behaviour.

Emergence of GRs was most frequent within Myrmicinae (39 orthogroups) than in ants (26 orthogroups). In contrast to ORs, the origin of GRs had no significant effect on the numbers of convergent losses in parasites (Fig. S2C).

### Parasites have substantially more genes under positive, relaxed and intensified selection

We next asked if the parasitic lifestyle might not only be characterised by a loss of genes but also be accompanied by significant changes in selection patterns.

Indeed we found that significantly more single-copy orthologs showed signatures of relaxed selection on the parasite branches (145 genes with an FDR *<* 0.05; 3.8% of tested genes) compared to host branches (36 genes, 0.9%; *χ*^2^=64.7; p=8.8×10*^−^*^16^; df=1; Fig. 4; Supplementary dataset 3). The genes under relaxed selection were enriched for GO-terms associated with protein modification and cell transport in parasites and response to stimulus in host species (Fig. 5). Interestingly, we also found significantly more genes under intensified selection (positive and purifying) in the parasite genomes (114 genes, 3.0%) than in the host genomes (81 genes, 2.1%; *χ*^2^=5.0; p=0.025; df=1; Fig. 4). Among the genes under intensified selection in parasites were some linked to lifestyle and multiple genes putatively involved in epigenetic regulation (Supplementary dataset 2c). In further support of our findings for convergent losses of genes involved in pathogen resistance, we found four immunity genes to be under relaxed selection in parasite lineages but only one in hosts (Supplementary dataset 3). One of these genes, neurocalcin, was under intensified selection in hosts. These results reflect the strategy of parasites to rely on the immunological armoury of their host to fend off diseases.

**Figure 4:**
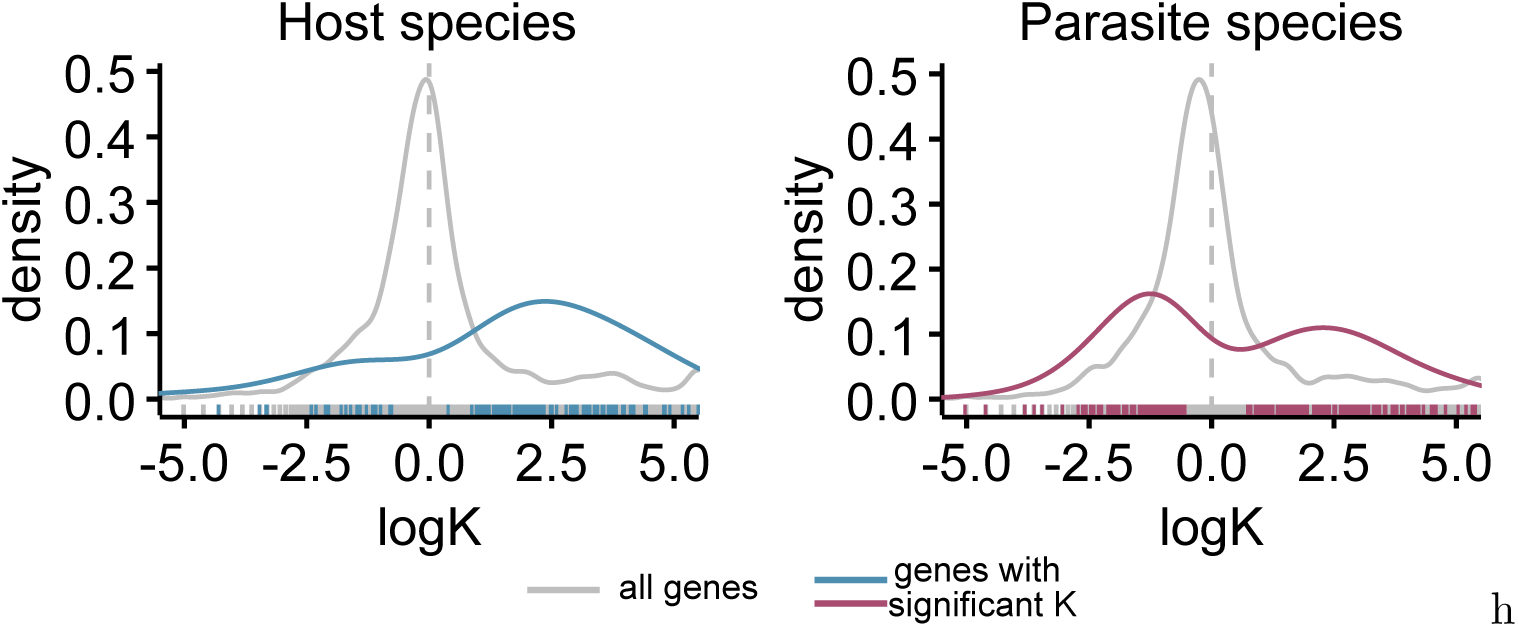
Relaxation and intensification of selection. Shown are the distributions of the selection intensity parameter (K), which represents the relative selection strength on the branches of interest (four terminal branches leading to the four parasite or host species) compared to the rest of the tree. Values of log(K) lower than 0 signify a relaxation, greater than 0 an intensification of selection. The grey curves show the distribution of log(K) among all genes, and the coloured curves show only genes with a significant K value (FDR *<* 0.05).

**Figure 5:**
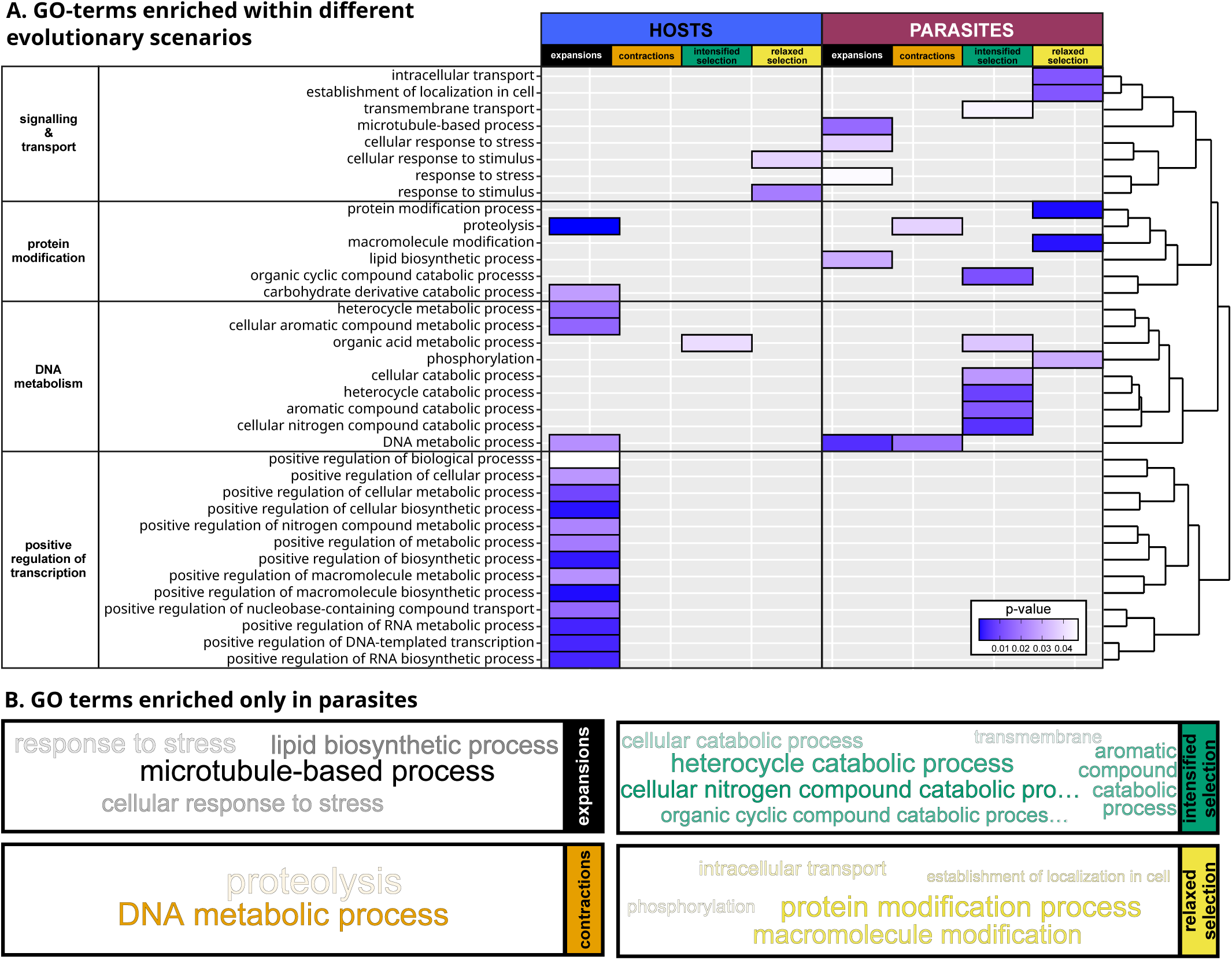
Functional enrichments. A. GO-terms that are enriched within significantly expanded and contracted gene families, as well as in genes under significant relaxation or intensification of selection. Genes under positive selection showed no enrichments and hence are not shown here. GO-terms are clustered by semantic similarity with colour intensity in the heatmap signifying the enrichment p-value. B. Tag-cloud of significant GO-terms that are enriched only in parasites and not in host species. Font size and colour intensity of the terms are related to the p-value (Supplementary dataset 4).

Furthermore, there were more genes with signatures of positive selection on the parasite branches (54 genes on single branches, 10 on two branches) compared to host branches (17 on single branches; *χ*^2^=26.4; p=2.8×10*^−^*^7^; df=1). The fact that most signatures of positive selection are found on single, maximally two branches indicates that there is no convergent selection on specific genes within host or parasite species. There were no enriched functions associated with genes under positive selection in hosts or parasites.

We next investigated the overlap between the different selection regimes and lifestyles (Fig. 6). We found the largest and most significant overlap between genes that were both under positive and intensified selection in parasites (25 genes, FE = 13.4; p = 4.3×10−23; FE: fold enrichment, obs:exp; Fig. 6). This indicates that the majority of the genes under significantly intensified selection are due to adaptation to the parasitic lifestyle. Given that our results showed divergent evolutionary trajectories between hosts and parasites, we were especially interested in genes that were under relaxed selection in parasites and under intensified and/or positive selection in hosts, and vice versa. Accordingly, we found significant overlaps in genes under relaxed selection in parasites but under intensified selection in hosts (9 genes; FE = 3.0; p = 2.8×10−3; Fig. 6, second stack from the left), under positive selection in hosts (3 genes; FE = 4.8; p = 0.023; Fig. 6, seventh stack), or both positive and intensified selection in hosts (1 gene; FE = 76.6; p = 0.013; Fig. 6, 13th stack). These 11 genes under relaxed selection in parasites but under positive or intensified selection in hosts covered varied functions including cell differentiation, neuronal genes and a member of the citric cycl,e ribosomal and mitochondrial genes (Supplementary dataset 2d).

**Figure 6:**
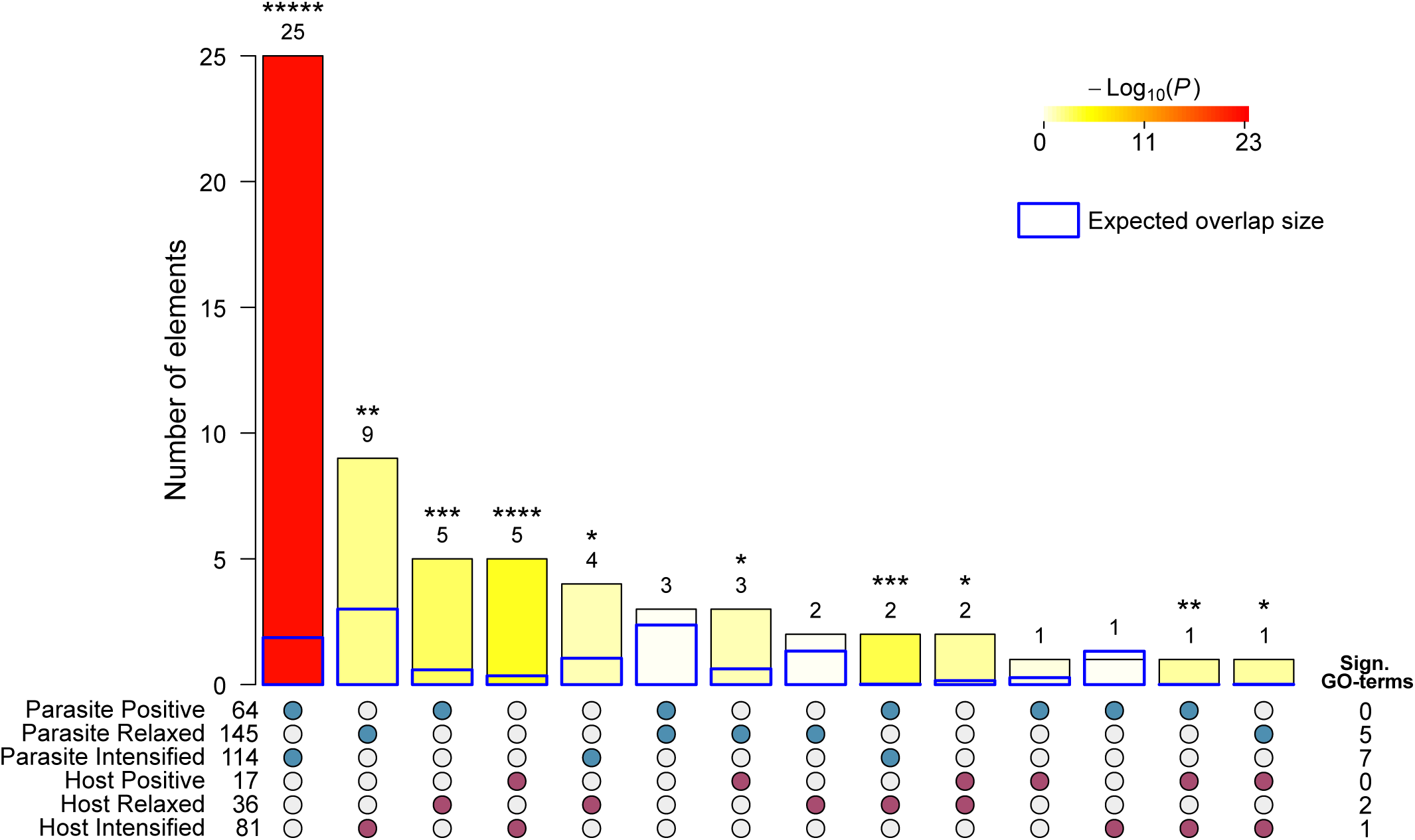
UpSet plot depicting the size and significance of gene overlaps between different selection regimes and species groups, the numbers of genes per category, as well as the number of significantly enriched GO functions per category. ***** p *<* 10*^−^*^5^; **** p *<* 10*^−^*^4^; *** p *<* 10*^−^*^3^; ** p *<* 0.01; * p *<* 0.05.

These genes might be important for brood care or foraging behaviours which have been lost in the parasites and are outsourced to the hosts. We found seven genes to be under relaxed selection in hosts but under intensified or positive selection in parasite species, indicating unique adaptations of the parasites (Supplementary dataset 2e).

### Putative *de novo* genes are consistently more frequent in parasite genomes

The total number of gene families in each genome ranges from 10,627 (*M. pharaonis*) to 14,788 (*T. ambiguus*) in our core set. An exception, reported already in [Schrader et al., 2021], is *C. obscurior*, which has a surprisingly high amount of 18,500 species-specific gene families (Fig. 1).

The total number of gene families that are specific to a single species (i.e. they are “private”, but may occasionally also contain a duplicate (paralog) in the same genome, see also Fig. S1) ranges from 393 (*T. longispinosus*) to 1079 (*A. ferox*), except for *C. obscurior*, which contains an exceptional amount of 5728 private gene families. Most of the species-specific gene families trace back to an older origin, while few of them emerged *de novo* in any of the genomes of the core data set.

For example, in *M. pharaonis*, among the 504 species-specific gene families, only one emerged likely *de novo*. As a less extreme example, in *H. sublaevis*, 119 (ca. 20%) out of 592 species-specific gene families emerged likely *de novo* (Fig. 1).

Between species, the number of putative *de novo* genes ranged from 54 in *T. longispinosus* to 125 in *H. sublaevis*.Parasites have a higher proportion of putative *de novo* genes than hosts do (phylogenetic ANOVA with “holm” postHoc test: t=-2.2036, p-value=0.033), but there were no significant differences in total numbers between non-hosts and parasites, and non-hosts and hosts (Fig. 7a).

**Figure 7:**
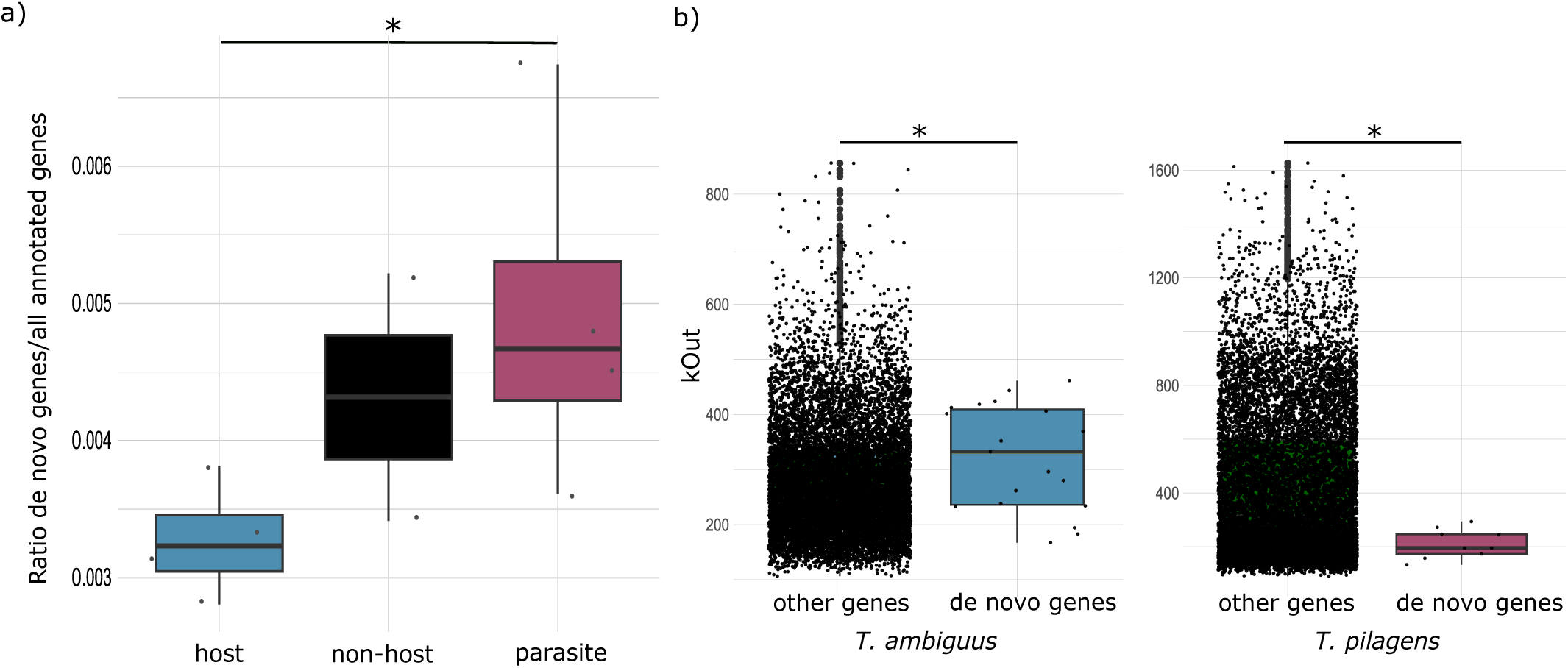
Prevalence and roles of putative *de novo* genes. a) The ratio of putative *de novo* genes differs across lifestyles. These results are based on the “core set” of 10 species. b) The connectivity of putative *de novo* genes outside of modules in comparison to all other genes is larger in *T. ambiguus* and smaller in *T. pilagens*.

**Figure 8:**
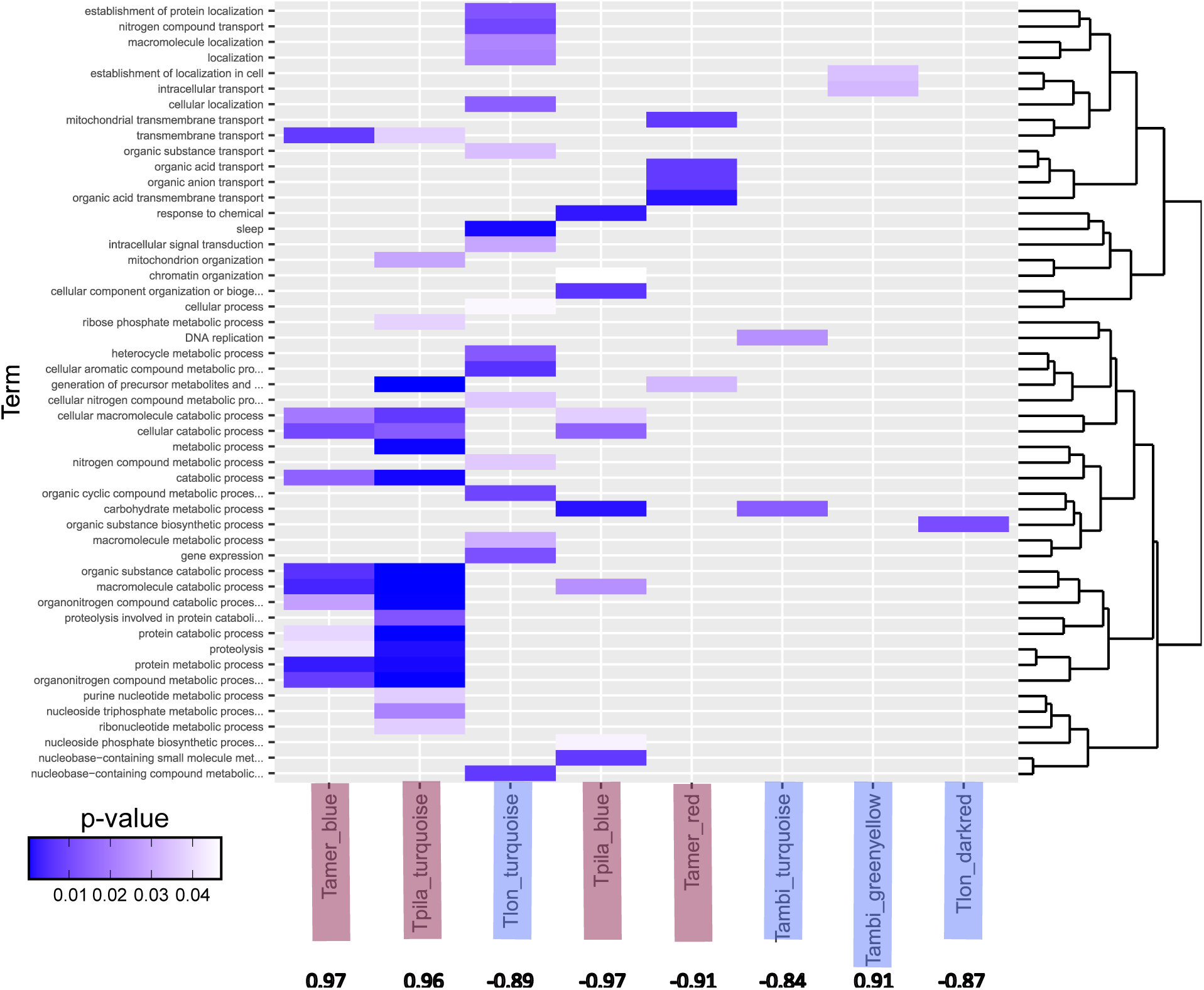
Enriched GO-terms within modules most significantly related to raids (up- and down-regulated) within two host and two parasite co-expression networks. Species in blue boxes are hosts and in red boxes are parasites. Numbers below each column represent correlation coefficients of module eigengene and raid behaviour. Enriched GO-terms (rows) are clustered by semantic similarity and WGCNA [Langfelder and Horvath, 2008] modules (columns) are also clustered by the similarity of p-values among the GO-terms.

To determine whether putative *de novo* genes are involved in raiding and nest-defence behaviour, or leading nest mates to the host nest (tandem running), we made use of data from two previous studies. We used these data sets to test for the conservation of gene-coexpression modules between hosts and parasites, and we investigated whether putative *de novo* genes were clustered in behaviour-associated modules.

The first study contrasted parasite raiding versus inactive behaviour [Alleman et al., 2018] plus host nest-defence versus foraging behaviour of three host-parasite pairs, two of which are included here (*T. longispinosus*-*T. americanus* and *T. ambiguus*-*T. pilagens*).

The second study investigated scouting, leading a tandem run versus following a tandem run in *T. americanus* and *T. longispinosus* [Alleman et al., 2019]. We found that *de novo* genes were distributed randomly across modules and did not cluster in specific, behaviour-associated modules (Bootstrap, number of modules 1000x: 0.05% *< x <* 0.95%). Moreover, the connectivity of *de novo* genes, as a measure of their “importance” in the raiding context, in comparison to other genes did not differ within gene co-expression modules in any of the species. In the host species *T. ambiguus*, however, the connectivity of *de novo* genes was higher, while in the parasite species *T. pilagens* they had lower overall connectivity outside of modules in the raiding context (Fig. 7b and Table S4a). There was no connectivity difference in the tandem-running context. These results, together with their generally relaxed selection regime and the fact that they have not yet become essential by being strongly integrated into the cellular networks, indicate that putative *de novo* genes are mainly retained by chance.

The only species where two putative *de novo* genes were differentially expressed was the host species *T. ambiguus*. Both *de novo* genes were up-regulated during the raid thus nest defence. *T. ambiguus* is also the only species where putative *de novo* genes had a trend towards higher expression levels in comparison to the other genes (t-test: t = −1.84, df = 21.023, p-value = 0.079). In all other species putative *de novo* genes had a generally low expression level and were not differentially expressed (Table S4b). Again the low and even expression levels of putative *de novo* genes indicate that most of them are retained by chance and non-essential, which is in keeping with a large number of recent studies on *de novo* genes [Schmitz et al., 2020, Grandchamp et al., 2024].

### Gene regulatory networks indicate common ancestry of recruitment behaviour

Differentially expressed genes (DEGs) had higher connectivity within and outside of their specific modules than genes that were not differentially expressed in the raiding as well as tandem-running context in all species (Table S4c).

In both host species from the raiding dataset (*T. longispinosus* and *T. ambiguus*) genes up-regulated during nest-defence had higher within-module connectivity than down-regulated genes. In parasites (*T. americanus* and *T. pilagens*) the opposite was observed, i.e. up-regulated genes during a raid had lower connectivity than down-regulated ones (Table S4d; Fig. S3). In all four species, genes that were not differentially expressed had the lowest connectivity, except in *T. longispinosus*, where genes up-regulated outside of the raiding context were similarly low. The lower connectivity of genes co-expressed during raiding in parasites, together with the peripheral network involvement of putative *de novo* genes, suggests that parasites employ more recently evolved and less essential genes whereas hosts co-express more highly connected and presumably evolutionary older modules.

Modules correlated with nest-defence in host species had very few enriched terms. In the two parasites (*T. americanus* and *T. pilagens*), the two modules most positively correlated with raids contained similar enriched GO terms, which indicates possible convergence in functions associated with raiding behaviour in these two species (Supplementary dataset 5). Many (≈30%) of these functions were related to catabolic processes which are involved in releasing energy [Russell and Cook, 1995].

Concerning similarities in gene co-expression, there was very strong gene module preservation between the host *T. longispinosus* and its parasite *T. americanus* in the tandem-running context, i.e., recruitment to new resources. Overall, the preservation score of gene modules associated with tandem-running was higher between parasites and hosts (mean=5.11) than in the raiding versus nest-defence scenario (t = −4.1218, df = 96.225, p-value = 7.97e-05; Fig. S4), indicating a common ancestry of tandem-running as recruitment strategy. The lower preservation between raiding modules corroborates the independent origins of the social parasites and their behaviours. However, we cannot rule out that the lower preservation may also stem from putatively more noisy whole-body transcriptomes analysed in the raiding study in comparison to brain transcriptomes in the tandem-running study.

### Genome-wide DNA methylation is similar in hosts and parasites

We next asked if epigenetic marks might underly the different expression profiles between hosts and parasites. Methylation levels were estimated *in silico* by comparing observed to expected counts of CpGs within coding regions of genes, consisting of exons, introns and 1kb flanking regions. A lower than expected CpG count indicates increased methylation levels. Overall, CpG depletion levels were relatively low, with 2.4 to 9.1% of genes with a CpG*_obs/exp_* less than 1 (0.4-1.2% with CpG*_obs/exp_ <* 0.75). CpG*_obs/exp_* did not differ between host (median: 1.36-1.39) and parasite species (median: 1.35-1.43). We analysed the function of methylated genes by concentrating on the genes within a CpG*_obs/exp_* lower than the bottom 10th percentile. A comparison of methylation patterns between hosts and their parasites revealed that at least half of the methylated genes were shared within host-parasite pairs (Fig. S5). In all cases, methylated genes were enriched for functions involved in various cellular and metabolic processes, methylation, DNA- and RNA-associated processes, thus metabolic and regulatory fundamental functions. A similar picture emerged when comparing methylated genes between lifestyles. There were many more shared genes (891 out of 4.651) and fewer lifestyle-specific genes (between 494-2.038; Fig. S6; Table S5) than expected by chance. Functions overrepresented in methylated genes in all three lifestyles mainly belonged to cellular- and metabolic-associated processes (Supplementary dataset 6; Fig. S6). This finding corroborates studies indicating that in ants mainly housekeeping genes are methylated, whereas genes responsible for phenotypic plasticity (also involved in caste differentiation) in ants are rather regulated via other epigenetic mechanisms, such as histone acetylation [Simola et al., 2016, 2013, Libbrecht et al., 2016, Kohlmeier et al., 2023].

## DISCUSSION

Parasites, by definition, exploit their host one way or another. Therefore, they are no longer in need of producing or expressing the exploited trait themselves. As the underlying gene repertoire may no longer be needed, evolutionary signatures may shift from purifying to relaxed selection at associated loci [Smith et al., 2015], and ultimately may get lost, leading to genome reduction [Tsai et al., 2013]. There are multiple examples of trait loss in multiple species [Wernegreen, 2005, Barrett et al., 2018, Oakeson et al., 2014, Niemiller et al., 2013]. However, the underlying genomic changes are largely under-explored. Here we investigated the genomic signatures of four independently evolved host-parasite pairs. This setting allows us to draw more general conclusions, in particular on the possible benefits of gene losses, than would be possible when considering one host-parasite pair alone. *Temnothorax* and *Harpagoxenus* parasites exploit the social behaviour of their hosts and keep host workers to perform the worker chores. At the same time, they evolved new behaviours to attack host colonies and steal the brood. We therefore expected to find numerous gene losses associated with worker behaviour but also the evolution of novel genes, or neofunctionalisation of existing genes associated with raiding behaviour.

We find that the evolution of existing gene families is more dynamic in hosts than in parasites since hosts show more species-specific expansions and contractions of gene families. We also find more convergent expansions of gene families in hosts than in parasites, but consistent with our hypothesis, we find more convergent contractions of gene families in the latter. Most strikingly, one third of ORs were lost in parasite genomes. Interestingly, many of these lost genes originated at the base of the ants, i.e., at or near the origins of sociality. Their loss in socially parasitic species may indicate that these genes were involved in the evolution of worker behaviour and complex social structures. This corroborates the results of another recent study showing differential expression of OR gene gene families lost in dulotic parasites in the antennae of *T. longispinosus* foragers and nurses [Caminer et al., 2023].

As social parasite workers do not perform typical worker tasks, we expected to find genes under relaxed selection, but at the same time also genes under positive selection putatively associated with the raiding behaviour. Indeed, we detected about twice as many genes under relaxed selection in parasites than in hosts. Moreover, we identified five times as many genes under positive selection in the parasites compared to hosts. This was unexpected since previous studies on social parasites in ants have either not detected relaxed or positive selection [Smith et al., 2015], or detected relaxed selection only [Schrader et al., 2021]. As these previous studies were conducted on inquiline species, which completely lost the worker caste, one may conclude that the loss of the worker caste may only lead to relaxed selection in genes associated with worker traits, if at all. In our species set, however, the worker caste was retained but shifted their behavioural profile to raiding, and these differences might be encoded in some of the positively selected genes.

Along those lines, we also found that parasite species had more putative *de novo* genes than host species. *De novo* genes have been shown to be frequent in ants [Wissler et al., 2013], but also to be frequently lost shortly after their emergence [Grandchamp et al., 2024, Gubala et al., 2017], to drive protein-novelty in *Drosophila* [Heames et al., 2020] and generally to evolve neutrally [Bornberg-Bauer et al., 2021, Schmitz et al., 2018]. Although frequent in parasites, putative *de novo* genes showed no association with the raiding phenotype. They had a generally low expression level, which is often observed in *de novo* genes [**?**], they were randomly distributed across various gene-co-expression modules, and their connectivity was comparable to most other genes.

However, two gene-coexpression modules, which were associated with the raiding phenotype, overlap to a large part in enriched functions in two parasite species, indicating convergence at the functional level.

### Conclusion

The evolution of eusociality is associated with the expansion of some gene families, which allowed social Hymenoptera to develop highly differentiated caste phenotypes, to communicate, and to evolve novel behaviours and morphologies. Further on in evolution, their social behaviours let to the emergence of social parasites that exploit their altruistic behaviours. As parasites exploit traits of their hosts, they do not need to express these traits anymore and the underlying genes may erode and get lost over time. At the same time, parasites may also express novel behaviours and traits associated with host exploitation, as is the case for social parasites that evolved raiding behaviours and morphological adaptations associated with these. Indeed, our results show that parasites evolved more putative *de novo* genes, in contrast to duplicated genes, the host species did, and exhibited more genes under positive selection. Furthermore, there were more gene losses and gene family contractions in the parasites than in the hosts. One prominent gene family that is strongly contracted in parasites is the OR-gene family and we show that the genes that are lost in parasites are often those, which emerged in ants on the evolutionary path to eusociality. Many of those have also been shown to be expressed differentially in workers conducting different tasks [Caminer et al., 2023]. In our study, we find evidence for convergent gene loss across parasitic species, species-specific gene losses and gains, as well as evidence for relaxed, but also positive selection. We conclude that relaxation and gene losses may be indicative of the loss of some worker behaviours, and *de novo* genes as well as positive selection signatures could be associated with the emergence of novel parasite traits such as raiding behaviour.

## METHODS

### Sample collection and genome sequencing

In a former study [Jongepier et al., 2022] we generated genomic data across three independent origins of social parasitism (formerly known as slave-making) in ants, of three parasite ant species (*Harpagoxenus sublaevis*, *Temnothorax ravouxi*, *T. americanus*), three host species (*Leptothorax acervorum*, *T. unifasciatus*, *T. longispinosus*) and two non-host outgroup species (*T. rugatulus*, *T. nylanderi*). Here we add one additional parasite-host pair, *T. pilagens*-*T. ambiguus*, as a fourth independent origin. These 10 species form the ***core set*** of analysed genomes in this study. Colonies were collected in the USA and Central Europe (Table S1). Multiple pupae from a single colony were pooled for each species to meet the requirements for whole-genome sequencing. For each species, two samples were prepared for WGS: 1) Pacbio Sequel long read library and 2) Illumina paired-end library. Selected host colonies were unparasitised, free-living colonies consisting of only a single species: the main host. In the parasite colonies, all brood should in principle be parasite brood because host workers do not produce offspring. Nonetheless, to rule out that any of the samples taken from parasite colonies are actually host brood (*e.g.* relics from previous so-called “slave raids”), only pupae of castes that are morphologically distinct from their host were selected for sequencing (*i.e.* queen pupae for *T. ravouxi*, and queens and worker pupae for *H. sublaevis*, *T. americanus*, and *T. pilagens*). Depending on the availability of pupae for each species and the need to pool DNA extractions due to low DNA content, between 28 and 156 pupal samples were obtained for WGS. We furthermore extracted RNA for genome annotation from one nurse, forager, queen and brood where available and pooled these for sequencing. Library preparation and sequencing were performed by Novogene under the umbrella of the Global Ant Genome Alliance [Boomsma et al., 2017]. For read statistics see (Table S2).

### Genome assemblies and polishing

Several genome assembly strategies were explored and assessed.For the final assembly, raw PacBio reads were assembled using the Canu [Koren et al., 2017] pipeline (parameter settings: correctedErrorRate=0.15), with K-mer based genome size estimates. The assemblies were polished in multiple rounds. First with Bowtie2aligned Illumina short reads (v2.3.4.1; Langmead and Salzberg, 2012) using Pilon (v1.22; parameter settings: diploid, fix=all; Walker et al., 2014). Second, raw PacBio reads were mapped against the assemblies with Minimap2 (v2.1; settings:-ax map-pb; Li, 2018), and then processed with Purge Haplotigs (Roach et al., 2018) and FinisherSC (in “fast” and “large” mode; Lam et al., 2015), followed by a final round of polishing with Pilon, Arrow (VariantCaller v2.1.0) and again Pilon. Assembly contiguity and completeness were assessed with QUAST (v3.1; Gurevich et al., 2013) and BUSCO (v3.0.2; Simão et al., 2015), respectively.

### Genome size estimates

Genome sizes were estimated using the following two strategies: 1) The K-mer distribution of the Illumina libraries, for which we used the KmerCountExact utility of BBMap [Bushnell, 2015]. This analysis was run for K-mer sizes ranging from 31 to 131, selecting the largest genome size estimate as input for the CANU [Koren et al., 2017] genome assemblies. 2) The coverage of the PacBio-based assembly, where we mapped the original PacBio reads back to the genome assembly using Minimap2 (v2.1; settings:-ax map-pb; Li, 2018) and determined the coverage frequency distribution with the readhist module of Purge Haplotigs [Roach et al., 2018]. The latter method is likely to yield higher and more correct genome size estimates than the former because large repetitive sequences are collapsed in the K-mer based estimate when they exceed Illumina read length. The coverage based genome size estimates of the final assemblies are very similar to the average genome size of Myrmicinae, which is 329.1 Mb [Tsutsui et al., 2008].

### Genome annotation

For whole genome annotation, raw Illumina reads were trimmed with Trimmomatic (v0.38; ILLUMINA-CLIP=2:30:10 LEADING=3 TRAILING=3 SLIDINGWINDOW=4:30 CROP=125 HEADCROP=18; [Bolger et al., 2014]. Trimmed reads were mapped to the genome using HiSat2 (v2.1.0; [Zhang et al., 2021], and the genome-guided transcriptomes assembled and merged with StringTie (v1.3.4; [Pertea et al., 2015]. Open reading frames were predicted using TransDecoder (v5.3.0; http://transdecoder.github.io/). Species-specific repeat libraries were created *de novo* using RepeatModeler (v1.0.11; [Flynn et al., 2020], LTRharvest [Ellinghaus et al., 2008] which is part of GenomeTools (v1.5.10; github.com/genometools/genometools) and TransposonPSI (https://github.com/NBISweden/TransposonPSI). The resulting libraries were merged and combined with the SINEbank (Insecta) [Vassetzky and Kramerov, 2013] repeat database and subsequently filtered for redundancies using CD-hit-est (v4.6.8; [Li and Godzik, 2006]), as well as for true proteins by blasting against the previously assembled transcriptomes. The repeats contained in the merged repeat library were classified with RepeatClassifier implemented in RepeatModeler, and then combined with the RepeatMasker repeat database before soft-masking the target genomes. To predict *ab initio* gene models, we used Braker (v2.1.6;[Hoff et al., 2019]), GeMoMa [Keilwagen et al., 2019] for homology-based gene prediction based on annotation information of seven other myrmicines, and PASA (https://github.com/PASApipeline/) RNA-seq-based gene prediction. We used EVidenceModeler [Haas et al., 2008] to combine the three types of evidence with the following weights: GeneMark and Augustus: 1; GeMoMa: 5; PASA:10. Chemoreceptor annotations were taken from [Jongepier et al., 2022].

### Outgroup genomes

For selection and gene family evolution analyses, the core set of 10 genomes were complemented by five outgroup genomes. *Pristomyrmex punctatus*, *Myrmecina graminicola*, and *Acanthomyrmex ferox* genome sequences and annotations were provided by the GAGA project [Boomsma et al., 2017]. These were supplemented by genome sequences and annotations of *Monomorium pharaonis* (GCF 013373865.1) and *Cardiocondyla obscurior* (Genome: GCA 019399895.1; annotations: [Errbii et al., 2024]), which were obtained from NCBI (accessed March 2023).

For the analysis of odorant receptors, we added the refseq assemblies and annotations of the 11 following genomes (NCBI, accessed August 2023):

*Odontomachus brunneus* (GCF 010583005.1), *Harpegnathos saltator* (GCF 003227715.2),

*Dinoponera quadriceps* (GCF 001313825.1), *Eciton burchellii* (GCA 020341155.1),

*Linepithema humile* (GCF 000217595.1), *Camponotus floridanus* (GCF 003227725.1),

*Formica selysi* (GCA 009859135.1), *Lasius fuliginosus* (GCA 949152525.1),

*Vespula vulagaris* (GCF 905475345.1), *Nasonia vitripennis* (GCF 009193385.2), and *Athalia rosae* (GCF 917208135.1).

### Annotation and analysis of chemoreceptors

Odorant receptors (ORs) and gustatory receptors (GRs) were annotated in the 10 core genomes and 11 outgroup genomes with Bitacora, v. 1.4 [Vizueta Moraga et al., 2020] at default settings. Bitacora was combined with blastp (v. 2.12.0+ [Camacho et al., 2009]), hmmscan (HMMER v. 3.3, [Potter et al., 2018]), and GeMoMa (v. 1.7.1, [Keilwagen et al., 2019]), and using a database of OR and GR sequences previously annotated in *Temnothorax* genomes [Jongepier et al., 2022].

Annotated sequences were filtered for partial hits with the Bitacora perl script get genes partial pseudo.pl, setting a minimum length of 220 amino acids for ORs and 300 amino acids for GRs. The presence of the correct PFAM domain (OR: PF02949; GR: PF08395) was checked in all annotated sequences, by running pfamscan, v. 1.6 [Finn et al., 2015], and annotations were reassigned if necessary. If gene annotations contained two domains, annotations were split based on alignments of the OR or GR database against the specific genomic region using exonerate (–model protein2genome –percent 75) [Slater and Birney, 2005].

To investigate evolutionary origins of chemoreceptor losses, ORs and GRs were clustered into orthogroups using OrthoFinder [Emms and Kelly, 2015]. We aimed to obtain a large number of orthogroups containing few gene copies per species, in order to detect as many losses of orthologs as possible. For this the MCL inflation factor (default: 1.5) in the OrthoFinder analysis was raised to 20. Consequently, for each orthogroup we analysed gene losses by checking for the existence of at least one gene copy within the genomes of each of the four host and four parasite species. Absence of a gene within any of the eight genomes was counted as a loss. Furthermore, OR and GR orthogroups were categorised according to their evolutionary origin based on the existence of gene copies:

(1) only within myrmicine species but not in any of the other analysed hymenopteran genomes (Myrmicine-CRs);
(2) outside of Myrmicinae but not outside of ants (Ant-CRs); (3) outside of ants (Hymenopteran-ORs)

To test for the enrichment of chemoreceptor losses in parasite or host lineages depending on their evolutionary origin, we carried out the following GLM from the lme4 R-package [Bates et al., 2015], assuming a poisson distribution: parasite losses ∼ evolutionary origin x host losses. Here, the variables ‘parasite losses’ and ‘host losses’ contained an integer from 0 to 4 for each orthogroup to represent the number of species, in which this orthogroup was lost. Evolutionary origin was a categorical variable representing the three aforementioned origins: Myrmicine, Ant, Hymenopteran. The results were plotted with interact plot from the R-package interactions [Long, 2019].

### Orthology clustering

Orthology was analysed for the 10 core set and 5 outgroup genomes. Protein sequences were extracted from the genomes based on GFF annotations using gffread [Pertea and Pertea, 2020] at default settings. Proteomes were cleaned by retaining only the longest isoform per gene locus and removing sequences with premature stop codons, using domain-world helper scripts (https://domain-world.zivgitlabpages.uni-muenster.de/dw-helper/index.html). Additional, putative transposable elements were detected and removed prior to orthology clustering with two methods: 1. using TransposonPSI (https://github.com/NBISweden/TransposonPSI); 2: when containing PFAM domains related to transposable elements (taken from [Min and Choi, 2019]) across all 15 ant species. For these proteomes orthologous groups were determined with OrthoFinder, v. 2.5.4, [Emms and Kelly, 2015] at default settings. Orthogroups are also referred to as “Gene families”, because both orthologs and paralogs of a gene family are pooled together.

### Functional annotations

PFAM domains and corresponding GO terms were annotated on the proteomes. The filtered proteomes of the 10 core and 5 outgroup species were annotated with Pfam domains in version 30.0 [Finn et al., 2015] with PfamScan v1.6 [Finn et al., 2016]. Gene Ontology (GO) terms were mapped to the PFAM annotations using pfam2go [Mitchell et al., 2015].

Putative functions were determined by searching for reciprocal best blast hits against the *Apis mellifera* genome. For this the current genome and gene annotations of *A. mellifera* were downloaded from NCBI (version: GCF 003254395.2; accessed January 2024). Protein sequences were extracted for *A. mellifera* as described above for the other analysed genomes. The proteomes of each of the 15 analysed genomes (core set plus outgroups) were blasted against the *A. mellifera* proteome and vice versa with blastp, v. 2.12.0+, [Altschul et al., 1990] using the output format 6 and otherwise default settings. Best reciprocal blast hits, i.e. if top hit of A:B corresponds to top hit of B:A, were extracted from blast output files with a custom python script. Orthogroups, which were produced with OrthoFinder [Emms and Kelly, 2015], were annotated with these *A. mellifera* orthologs, PFAM domains, and GO-terms of all gene members within an orthogroup.

GO-term enrichment analyses were performed with the package topGO in R using the classic and weight algorithms [Alexa et al., 2010]. Significantly enriched GO-terms (p-value *<* 0.05) were grouped based on semantic similarity using Wang’s graph-based method from the R package GOSemSim [Yu et al., 2010] and visualized using the library ggplot2 for heatmaps [Wickham, 2016] and ggdendro for the dendrogram of GO terms [de Vries and Ripley, 2022].

### Gene family distribution and *de novo* gene detection

Python scripts using Biopython were used to search how the gene families generated with OrthoFinder were distributed in the tree. All orthogroups that contained at least one gene detected in the 15 species of the tree are referred to as “Gene families shared by all”. In such gene families, some species have several orthologous/paralogous genes. Indeed, for each branch, the figure shows whether the “Gene family shared by all” contains duplications in the species (”with duplications”) or not (”without duplications”). Orthogroups that contain genes shared by several species of the tree but not all are referred to as “Gene family shared by some”. If such gene families contain genes only detected in hosts, they are referred to as “Gene families only found in hosts”. On the contrary, when these gene families contain genes only detected in parasites, they are referred as “Gene families only found in parasites”. Orthogroups that contain genes found in only one species are referred as “Gene specific to a single species”. Among these orthogroups, some contain several genes from the species, and are referred to as “Gene specific to a single species in duplication”, while orthogroups containing only one gene are called “Gene specific to a single species found in singleton”. The total number of gene families ranged from 10,627 (*M. pharaonis*) to 14,788 (*T. ambiguus*), with the exception of *C. obscurior* which had a surprisingly high amount of species-specific genes families (18,500). The total number of gene families that were specific to a species range from 1079 (*A. ferox*) to 393 (*T. longispinosus*), again with Cobs standing out (5728 gene families). Among all species-specific gene families, all translated genes were set as query for protein BLAST (default parameters) to validate the lack of homology to any other genes in other species plus outgroup species. All species-specific gene families, i.e. without homology to outgroup proteins, were considered as *de novo* emerged. Non-coding homologs were searched for in syntenic regions of sister species (Supplemetary dataset 7).

### Identification of losses and gains

Gene families were identified and classified using the above mentioned orthogroups. Orthogroups and genes within were classified into various categories depending on whether they were shared by all species, a subset of species (e.g. hosts only, parasites only,…), to species-specific using an inhouse python script. To determine whether a gene family or a single gene classified as species-specific or emerged *de novo*, we ensured that the gene(family) did not show homology to any other gene in any other species in our data set. To determine which of the species-specific genes emerged *de novo* and which are suspected to have arisen via the fast evolution of existing genes, all species-specific genes were used as queries for a protein BLAST search against: 1. all proteins from all other species of the tree; 2. The proteome of 6 ants outgroup species: *Acromyrmex echinatior; Atta cephalotes; Camponotus floridanus; Monomorium pharaonis; Ooceraea biroi and Pogonomyrmex barbatus* (https://antgenomes.org/).

### Relaxed and positive selection analyses

For selection analyses, single copy ortholog groups (orthogroups with one gene copy in each of the 10 core and 5 outgroups genomes) were determined with OrthoFinder [Emms and Kelly, 2015] and nucleotide sequences were aligned with pal2nal [Suyama et al., 2006] based on protein alignments that were performed with prank, v.170427, with-F and-once settings [Loytynoja, 2013]. Nucleotide alignments were trimmed with gblocks (v.0.91b;-t=c-b5=h) [Castresana, 2000]. With these trimmed alignments of all single copy orthologs two selection analyses (RELAX and aBSREL) from the HyPhy suite [Pond and Muse, 2005] were performed. To determine signatures of relaxed and intensified selection on the four host or four parasite branches compared to the rest of the tree, we ran RELAX at default settings. K, the relaxation factor, and corresponding p values were extracted from output files. P-values were corrected for multiple testing with the FDR method. Selection on genes was determined to be significantly relaxed with a k *<* 1 and an FDR *<* 0.05 or intensified with a k *>* 1 and FDR *<* 0.05. To test for positive selection within single copy orthologs, aBSREL was run on the same alignments with the following additional settings: –multiple-hits Double+Triple –srv Yes. The first argument allows multiple simultaneous mutations (1, 2 or 3) within a codon, while the second argument allows substitution rates to vary among synonymous sites. Testing for significance in overlaps between genes under positive, relaxed and intensified selection was carried out with the SuperExaxtTest R package [Wang et al., 2022].

### Methylation estimations

We estimated methylation levels by comparing observed to expected numbers of C-G nucleotide pairs (CpG) within gene bodies. For this analysis, gene body constituted the full genomic sequence of a gene region, including 1kb of flanking regions. Expected CpG counts were calculated based on the product of the relative frequencies of Cs and Gs within a gene sequence. The observed frequency of CGs was then divided by the expected frequency to obtain CpG*_obs/exp_* values. DNA methylation at CpG sites is known to lead to a depletion in observed CpGs as the cytosines of methylated CpGs often mutate to thymines [Bewick et al., 2016]. A CpG*_obs/exp_ <* 1 is therefore an indication of increased levels of methylation.

### Gene network analyses

We used the WGCNA package [Langfelder and Horvath, 2008] to conduct weighted correlation network analyses in R based on transcriptome data from two studies [Alleman et al., 2018, 2019]. The function *pickSoftThreshold* was used to determine the best-fitting soft threshold power per data set, which turned out to be 12 and 11 respectively. We additionally tested for module trait correlation with “before raid” and “during raid” as trait for the [Alleman et al., 2018] data set, and “scout”, “leader” and “follower” to associate modules to these different behaviours from [Alleman et al., 2019]. We employed the function *intramodularConnectivity* to investigate the connectivity of genes within and outside modules. The package hdWGCNA was used to determine preserved modules between all possible species pairs of the six species included in the network analyses by using each species as the reference and vice versa. We used the online tool “Venn Diagrams” (https://www.biotools.fr/misc/venny) to construct Venn diagrams.

## Data availability statement

All raw DNA sequence data underlying this study as well as the novel genome assemblies are deposited in the National Centre for Biotechnological Information (NCBI) under BioSample accession number: PR-JNA750352.

## Author contributions

E.B.-B., J.H., and S.F. conceived the study and acquired funding. E.B.-B., J.H., S.F. and B.F. designed the study. E.J., A.S., C.F. and M.S. assembled and annotated the genomes. A.G. identified *de novo* genes. M.H. performed the gene family evolution and selection analyses and identified CpG islands. B.F. performed the gene network and preservation analyses. M.H., E.B.-B. and B.F. wrote the first draft of the paper which was revised by all authors who also contributed to interpretation of data. Authors declare no conflict of interest.

## Acknowledgements

We thank Jonas David Wittig for help with collecting *T. pilagens* and *T. ambiguus* colonies at the Sleeping Bear National Sand Dunes, Michigan, for which we had obtained a collection permit from park authorities. We thank Lukas Schrader and the Global Ant Genome Alliance (GAGA, Vizueta et al., submitted) for their help with sequencing the genomes of *Pristomyrmex punctatus*, *Myrmecina graminicola*, and *Acanthomyrmex ferox*. This work was supported by the Deutsche Forschungsgemeinschaft (Grant Nos. BO 2544/12-1 and BO 2544/20-1 to E.B.-B., FO 298/20-1 to S.F., and HE 1623/40-1 to J.H). E.B.-B. acknowledges that parts of this work was conducted while visiting the Okinawa Institute of Science and Technology (OIST) through the Theoretical Sciences Visiting Program (TSVP).

